# OTX2 signals from the choroid plexus to regulate adult neurogenesis

**DOI:** 10.1101/243659

**Authors:** Anabelle Planques, Vanessa Oliveira Moreira, Chantal Dubreuil, Alain Prochiantz, Ariel A Di Nardo

## Abstract

Proliferation and migration during adult neurogenesis are regulated by a microenvironment of signaling molecules originating from local vasculature, from cerebrospinal fluid produced by the choroid plexus, and from local supporting cells including astrocytes. Here, we focus on the function of OTX2 homeoprotein transcription factor in the mouse adult ventricular-subventricular zone (V-SVZ) which generates olfactory bulb neurons. We find that OTX2 secreted by choroid plexus is transferred to supporting cells of the V-SVZ and rostral migratory stream. Deletion of *Otx2* in choroid plexus affects neuroblast migration and reduces the number of olfactory bulb newborn neurons. Adult neurogenesis was also decreased by expressing secreted single-chain antibodies to sequester OTX2 in the cerebrospinal fluid, demonstrating the importance of non-cell autonomous OTX2. We show that OTX2 activity modifies extracellular matrix components and signaling molecules produced by supporting astrocytes. Thus, we reveal a multi-level and non-cell autonomous role of a homeoprotein and reinforce the choroid plexus and astrocytes as key niche compartments affecting adult neurogenesis.

**Significance Statement:** Cerebrospinal fluid, local vasculature and non-neurogenic astrocytes are niche compartments that provide a microenvironment for regulating adult mouse neurogenesis. We show that OTX2 homeoprotein secreted by choroid plexus into the cerebrospinal fluid is transferred into non-neurogenic astrocytes of the ventricular-subventricular zone and rostral migratory stream where it regulates extracellular matrix and signaling factors. This non-cell-autonomous activity impacts the number of newborn neurons that integrate the olfactory bulb. Thus, we reveal a multi-level role for OTX2 and reinforce the choroid plexus as a key niche compartment affecting adult neurogenesis.

## Introduction

Neurogenesis in the adult mouse brain provides continuous replacement of interneurons in olfactory bulbs (OB) and is important for olfaction-based learning (Lledo and Valley, 2016). Neural stem cells (NSCs), located in the adult ventricular-subventricular zone (V-SVZ) lining the lateral cerebral ventricles, give rise to progenitor cells (first transient amplifying cells that generate migrating neuroblasts) that migrate though the rostral migratory stream (RMS) to reach the OB where they differentiate as interneurons and integrate into local circuitry. This process is regulated not only by factors intrinsic to the NSCs and progenitors but also by their microenvironment consisting of factors from local vasculature, neuronal circuits, cerebrospinal fluid (CSF) and local supporting cells (Lehtinen et al., 2013; Silva-Vargas et al., 2013; Bjornsson et al., 2015). It has been shown that NSCs contact vascular endothelial cells and project into the ventricular wall where their cilia contact CSF, which participates in regulating proliferation and migration through mechanical influence and CSF-borne factors (Sawamoto et al., 2006; Mirzadeh et al., 2008; Shen et al., 2008; Petrik et al., 2018).

The CSF is produced within brain ventricles by the choroid plexuses (ChP) which express many secreted factors found to influence neurogenesis, including growth factors, morphogens and guidance clues (Falcão et al., 2012; Kokovay et al., 2012; Silva-Vargas et al., 2016). Examples of CSF-borne factors include IGF2, which promotes adult NSC renewal (Ziegler et al., 2015), IGF1 and BMP5, which promote NSC colonies (Silva-Vargas et al., 2016), NT-3, which promotes quiescence (Delgado et al., 2014), and SLIT2, a chemorepulsive factor that participates in neuroblast migration (Nguyen-Ba-Charvet et al., 2004; Sawamoto et al., 2006). Neuroblasts and NSCs also contact local non-neurogenic astrocytes (Lois et al., 1996; Shen et al., 2008) which secrete factors that help establish and maintain the microenvironment and can influence neurogenesis (reviewed in Gengatharan et al., 2016). Such factors include DLK1, which induces NSC self-renewal (Ferrón et al., 2011), and thrombospondin 4 (THBS4), which guides neuroblast migration (Girard et al., 2014).

Homeoproteins are key regulators of neurogenesis both during embryogenesis and in the adult (Curto et al., 2014; Prochiantz and Di Nardo, 2015). This class of transcription factors have the property to act both cell-autonomously and non-cell-autonomously after secretion and internalization in target cells. The Otx2 homeoprotein is a regulator of embryonic development and embryonic neurogenesis (Yang et al., 2014; Hoch et al., 2015; Acampora et al., 2016) but its role on adult neurogenesis has not been investigated. OTX2 is expressed by ChP, secreted in CSF, and transfers in brain parenchyma to regulate cortical plasticity in postnatal mice (Spatazza et al., 2013; Lee et al., 2017). We hypothesized that OTX2 in the CSF might also accumulate in adult neurogenic niches and impact neurogenesis. Indeed, we find V-SVZ neurogenesis is regulated by OTX2 expression and secretion by the CP, and find that OTX2 transfers into supporting astrocytes to control the expression of extracellular factors that impact neuroblast migration.

## Materials and Methods

### Animals

S129 mice were purchased from Charles River Laboratories. The *Otx2^lox/lox^* (Otx2-lox) mice (S129 background) were kindly donated by Dr. T. Lamonerie (Fossat et al., 2006) and *Otx2*^+/*GFP*^ mice were kindly donated by Dr. A. Simeone (Acampora et al., 2009). The single chain antibody (scFv) knock-in mouse model (scFv-Otx2 mice, C57BL/6J background) were previously described (Bernard et al., 2016). Adult mice (3-months old) of either sex were used in all experiments, with littermates distributed equally between control and treated groups while maintaining gender balance. All animal procedures, including housing, were carried out in accordance with the guidelines of the European Economic Community (2010/63/UE) and the French National Committee (2013/118). This research (project no. 00704.02) was approved by Ethics committee n° 59 of the French Ministry for Research and Higher Education. Mice were conventionally raised (12:12 hours light:dark cycle) in cages with red tunnels, nesting cotton, and with food and water *ad libitum*.

### Mouse surgery

For surgical procedures, animals were anesthetized with Xylazine (Rompun 2%, 5 mg/kg) and Ketamine (Imalgene 500, 80 mg/kg). Cre-Tat recombinant protein (8 to 20 μg/ml in 400 mM NaCl, 15% DMSO) or vehicle was injected as previously described (Spatazza et al., 2013). 10 days after Cre-Tat injection, mice were injected intraperitoneally with 150 mg/kg of BrdU (Sigma B9285 at 10 mg/ml diluted in NaCl 0.9%). For OB integration experiments, mice were injected with BrdU twice daily for five days, and then housed for 3 weeks prior to perfusion with PBS. For proliferation experiments, mice were injected with one pulse of BrdU and then perfused with PBS two hours later. Brains were cut in half along the anterior-posterior axis: the anterior half was fixed for 48 h at 4 °C in 4% formaldehyde-PBS, immersed overnight in 20% sucrose PBS, and processed for cryostat sectioning; the posterior half was used to recover 4^th^ ventricle ChP for quantitative PCR analysis of *Otx2* expression.

### Astrocyte cell culture

Primary astrocytes were prepared from cortex of newborn mouse (P0 to P4) of either sex and cultured in 10 mL of DMEM high glucose, 10% serum, 1x antimitotic/antibiotic (Même et al., 2006). After 7 days, astrocytes were passaged into 6-well plates at 400 000 cells/ml, incubated 14 days, and then were treated with 1% Ara-C (in DMEM high glucose, 1% B27, 1x antimitotic/antibiotic) for 24 h prior to treatment with 100 ng/ml OTX2 protein. After 24 h, treated astrocytes were processed for quantitative PCR analysis.

### Immunohistochemistry and histology

Frozen brains were sectioned (25 or 40 μm) by cryostat and stored at -20°C. For all staining experiments, slides were thawed 15 min at room temperature, hydrated in PBS, and incubated for antigen retrieval in a steamer for 15 min with 10 mM citrate buffer, pH 6. After cooling, sections were washed with PBS and incubated in blocking buffer (0.5% Triton-X, 10% normal donkey serum (Abcam) in PBS) for 30 min. Primary antibodies were incubated overnight at room temperature (1/100 rat anti-BrdU OBT0030 AbD Serotec; 1/400 rabbit anti-Caspase #9661 Cell Signaling; 1/250 rabbit anti-OTX2 ab92326 Abcam; 1/1000 guinea pig anti-DCX AB2253 Millipore; 1/200 mouse anti-GFAP G6171 Sigma; 1/1000 chicken anti-Vimentin AB5733 Millipore; 1/1000 chicken anti-GFP ab13970 Abcam). After PBS washes, secondary antibodies (1/2000 donkey anti-IgG Alexa Fluor, Invitrogen) were incubated 2 h at room temperature. For DCX labeling, sections were washed with PBS, incubated 2 h at room temperature with secondary antibody (1/500 biotinylated goat anti-guinea pig ab6907 Abcam), washed again with PBS, and incubated 2 h with conjugated streptavadin (1/2000 Alexa Fluor Streptavidin, Invitrogen). Sections were washed in PBS, dried and mounted in DAPI Fluoromount-G (Southern Biotech). TUNEL was performed with the *in situ* cell death detection kit (11684817910 Roche) according to manufacturer instructions. *In situ* hybridization was performed as previously described (Spatazza et al., 2013).

### Imaging and quantification

Imaging was performed with an SP5 inverted confocal microscope (Leica), a Nikon 90i upright widefield, or a Spinning-disk confocal with constant parameters. For BrdU analysis, one 40 μm coronal section every 200 μm was quantified. BrdU counting in the granule cell layer (GCL) and glomerular layer (GL) from the same mice was performed on 6 sections per animal from bregma +4.0 to its anterior end using an in-house ImageJ macro that automatically detects cells within an ROI. Cell number was normalized by the ROI area for each image; the mean for each animal was reported. This method was validated in initial experiments where counting by hand in parallel gave similar results. BrdU counting in the V-SVZ was done by hand on 8 x 40 μm coronal sections every 200 μm per animal, from bregma +1.34 to bregma -0.58mm, which represents the anterior V-SVZ in which proliferation is more sustained (Fiorelli et al., 2015). Cell numbers were normalized by ventricular wall length.

### Quantitative PCR

Total RNA from 4^th^ ventricle ChP was extracted with the RNeasy Lipid Tissue Mini Kit (Qiagen) with DNA removal. RNA and protein were extracted for ECM composition analysis using the AllPrep DNA/RNA/Protein Mini Kit (Qiagen). RNA was processed with the QuantiTect Reverse Transcription Kit (Qiagen). cDNA was diluted 1/10 with RNase-free water for quantitative PCR, samples, which were analyzed in triplicate with a LightCycler 480 II (Roche) and SYBR Green I Master mix. After *T_m_* profile validation, gene expression was determined by the 2^-*ΔΔCt*^ method with hypoxanthine guanine phosphoribosyl transferase (HPRT) as the control gene. For expression analysis, mean expression of vehicle-injected mice or control-treated cells was used for comparison.

### Microdissection

For biochemical analysis, mice were sacrificed by cervical elongation. Brain was placed in cold PBS, dorsal side facing up, and two equidistant longitudinal incisions in each OB were made to divide them in 3 parts while keeping them attached to the rest of the brain. The V-SVZ microdissection was performed as previously described (Mirzadeh et al., 2010). Afterwards, the hemispheres were placed once again dorsal side facing up and the two sagittal cuts starting from the OB incisions were pursued along the entire length of the brain. The central piece of tissue was laid on its side and RMS was excised as shown in Fig. 2*D*. All tissue samples were stored at -80°C.

**Figure 1.**
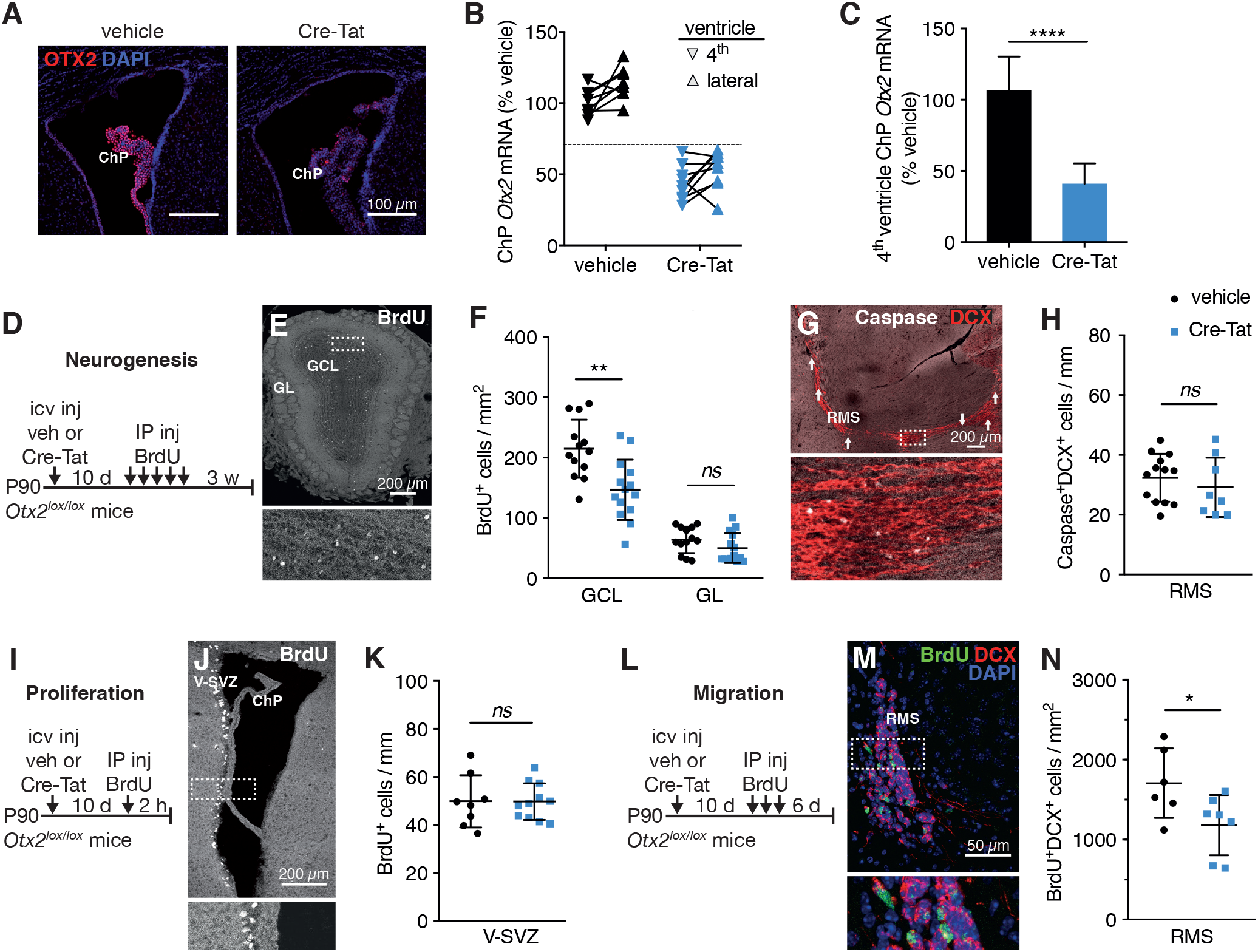
OTX2 knockdown in the choroid plexus reduces newborn neurons in the olfactory bulb. ***A-C***, Conditional *Otx2* knockdown with *Otx2^lox/lox^* mice. ***A***, Staining for OTX2 in lateral ventricle choroid plexus 35 d after icv injection of vehicle or Cre-Tat. ***B***, Comparison of *Otx2* expression levels between 4^th^ and lateral ventricles from the same brain (linked data points represent individual mouse) after vehicle or Cre-Tat icv injection. Dotted line represents 30% decrease in *Otx2* expression. ***C***, Quantitative PCR analysis of *Otx2* expression in 4^th^ ventricle (\*\*\*\**P*<0.0001, Mann-Whitney test, 5 experiments). ***D***, Schematic of adult (3 months) neurogenesis study paradigm. ***E***, Staining for BrdU in a coronal section of olfactory bulb 3 weeks after BrdU injections. ***F***, Quantification of BrdU-positive cells in GCL and GL of the olfactory bulb (\*\**P*<0.01, *ns*=*P*>0.05, t-test, 3 experiments). ***G***, Staining for caspase and DCX in a sagittal section of the RMS. Arrows indicate caspase-labeled neuroblasts throughout the entire RMS. ***H***, Quantification of caspase/DCX co-labeled cells in RMS (*ns*=*P*>0.05, t-test, 3 experiments). ***I***, Schematic of adult V-SVZ proliferation study paradigm. ***J***, Staining for BrdU in a coronal section of the V-SVZ 2 hours after BrdU injection. ***K***, Quantification of BrdU-positive cells in V-SVZ (*ns*=*P*>0.05, t-test, 2 experiments). ***L***, Schematic of adult migration study paradigm. ***M***, Staining for BrdU and DCX in a coronal section of the RMS. ***N***, Quantification of BrdU/DCX co-labeled cells in RMS (\**P*<0.05, t-test, 2 experiments). ChP, choroid plexus; GCL, granule cell layer; GL, glomerular layer; icv, intracerebroventricular; IP, intraperitoneal; RMS, rostral migratory stream; V-SVZ, ventricular-subventricular zone. Error bars represent SD.

**Figure 2.**
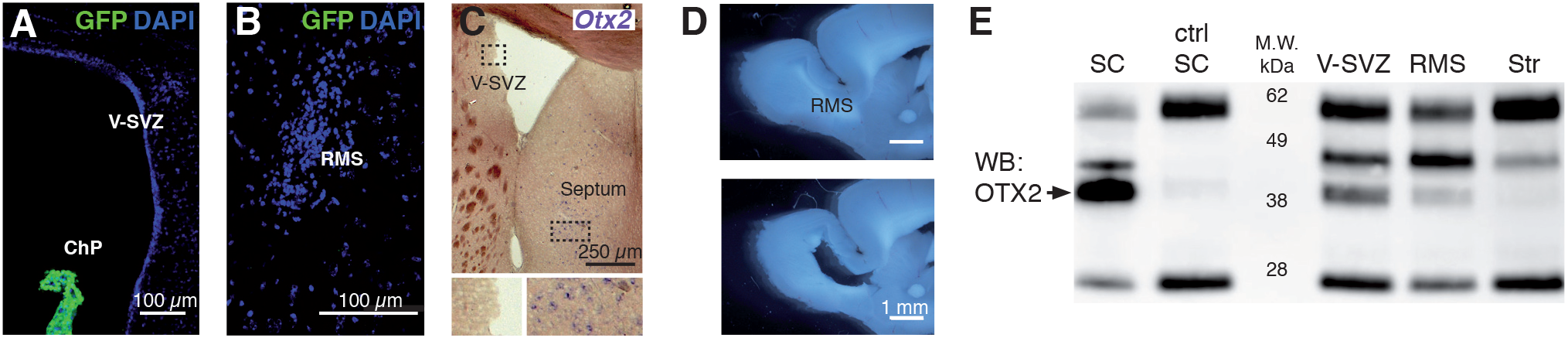
OTX2 in RMS and V-SVZ is non-cell-autonomous. ***A-C***, Absence of *Otx2* expression in V-SVZ (***A, C***) and RMS (***B***) as shown by GFP staining in coronal sections of *Otx2*^+/*GFP*^ mice (***A, B***) and by *in situ* hybridization for *Otx2* mRNA in coronal sections of wild-type mice (***C***). Note the expression of GFP in choroid plexus (***A***) and of *Otx2* in septum (***C***). ***D***, Representative sagittal section of RMS microdissection. ***E***, Western blot (WB) analysis of OTX2 immunoprecipitation from lysates of various brain regions (ctrl, IgG control). Bands include OTX2 (<40 kDa), IgG (25 and 55 kDa), and unknown (~45 kDa). ChP, choroid plexus; RMS, rostral migratory stream; SC, superior colliculus; Str, striatum; V-SVZ, ventricular-subventricular zone.

### Immunoprecipitation and Western blot

Microdissections were triturated in 100 μl of lysis buffer (150 mM NaCl, 1% Triton, 100 mM Tris, pH 7.4, 10 mM EDTA, 1X protease/phosphatase inhibitor (Roche)) per 10 mg of tissue and incubated 10 min on ice. Supernatant was recovered in low-bind tubes after centrifugation for 10 min at 16 000 g. Antibody was added (2.5 μg of anti-OTX2 (ab21990, Abcam) or anti-IgG (ab27478, Abcam) and incubated overnight at 4 °C with rotation. Protein A sepharose beads (GE Healthcare) were washed twice and resuspended 1:1 with lysis buffer. Beads (30 μl) were added to each sample and incubated 3 h at 4 °C. Beads were recovered by centrifugation for 1 min at 16 000 *g* and washed 5 times with 1 mL of cold wash buffer (Lysis buffer with 500 mM NaCl). Proteins were eluted from the beads with 30 μl of 2X laemmli buffer, heated 5 min at 95°C, and stored at -20°C.

Samples were separated on NuPage 4-12% Bis-Tris pre-cast gels (Invitrogen) with 1X MES buffer and antioxidant (Invitrogen) then processed for Western Blotting for OTX2 detection. Membranes were blocked in 5% non-fat milk, TBS, 0.2% Tween, incubated with anti-OTX2 (in-house mouse monoclonal) overnight at 4 °C then incubated with HRP-conjugated antimouse (GeneTex) for 2 h at room temperature. Chemiluminescence reaction was performed with ECL femto (Thermo Scientific). These immunoprecipitation experiments were performed 3 times with similar results each time.

### Statistical analysis

Prism 8 software (GraphPad) was used for all statistical analysis. Sample size was determined after preliminary experiments. Quantification was normalized by the mean value of vehicle-injected mice samples. Normal data distribution was assessed by the D’Agostino & Pearson omnibus normality test. If normal test was not passed, Mann-Whitney was applied; otherwise, unpaired t-test with Welch’s correction was used (noted t-test in Legends). See Figure Legends for relevant statistical tests.

## Results

### Otx2 knockdown in choroid plexus reduces newborn neurons in the olfactory bulb

Intracerebroventricular (icv) injections of vectorized Cre recombinase (Cre-Tat) leads to specific recombination of *Otx2* in lateral ventricle ChP of *Otx2^lox/lox^* (Otx2-lox) mice (Spatazza et al., 2013). With this paradigm, we find that Cre-Tat icv injection leads to a strong decrease in OTX2 protein in lateral ventricle ChP compared to vehicle-injected mice (Fig. 1*A*), and also a decrease in *Otx2* mRNA expression in both lateral and 4^th^ ventricle ChP (Fig. 1*B*). While the decrease is not necessarily proportional between both ChP, a decrease of at least 30% in the 4^th^ ventricle ChP reflects a decrease of at least 30% in the lateral ventricle ChP of the same mouse (Fig. 1*B*). Therefore, the 4^th^ ventricle ChP qPCR analysis is a valid method to assess *Otx2* expression levels in lateral ChP so that anterior tissues can be used for histological quantification. Otx2-lox mice with less than 30% *Otx2* expression knockdown in the 4^th^ ventricle after Cre-Tat icv injection where excluded from subsequent analysis. For the experiments described below, Cre-Tat injections in lateral cerebral ventricles lead to ~60% decrease in *Otx2* mRNA expression on average in the 4^th^ ventricle ChP (Fig. 1*C*).

To assess whether *Otx2* knockdown in ChP alters V-SVZ neurogenesis, we used BrdU treatments to measure either newborn neuron integration in OB (three weeks post-injection, Fig. 1*D-F*) or cell proliferation in V-SVZ (two hours post-injection, Fig. 1*I-K*) after Cre-Tat or vehicle icv injections. We found a significant decrease in the density of granule cell layer (GCL) newborn neurons but no change in the density of glomerular layer (GL) newborn neurons after *Otx2* knockdown (Fig. 1*F*). This change in density was not due to gross structural changes as no significant change in OB area was observed (Table 1) and was not dependent on gender (Table 1). Analysis of caspase-expressing cells revealed no change in cell death in the RMS of Cre-Tat-injected mice (Fig. 1*G,H*), suggesting that OTX2 does not affect neuroblast survival. However, *Otx2* knockdown in ChP does not alter V-SVZ cell proliferation (Fig. 1*K*), suggesting that the observed decrease in newborn neurons is due to effects on neuroblast migration. To test this hypothesis, we used a migration paradigm in which we evaluated the density of BrdU/DCX co-labeled cells in the RMS 6 days after final BrdU injection (Fig. 1*L-N*). We observed a significant decrease in Cre-Tat injected animals, suggesting that fewer neuroblasts are leaving the V-SVZ and/or that they are migrating slower (Fig. 1*N*).

**Table 1:**
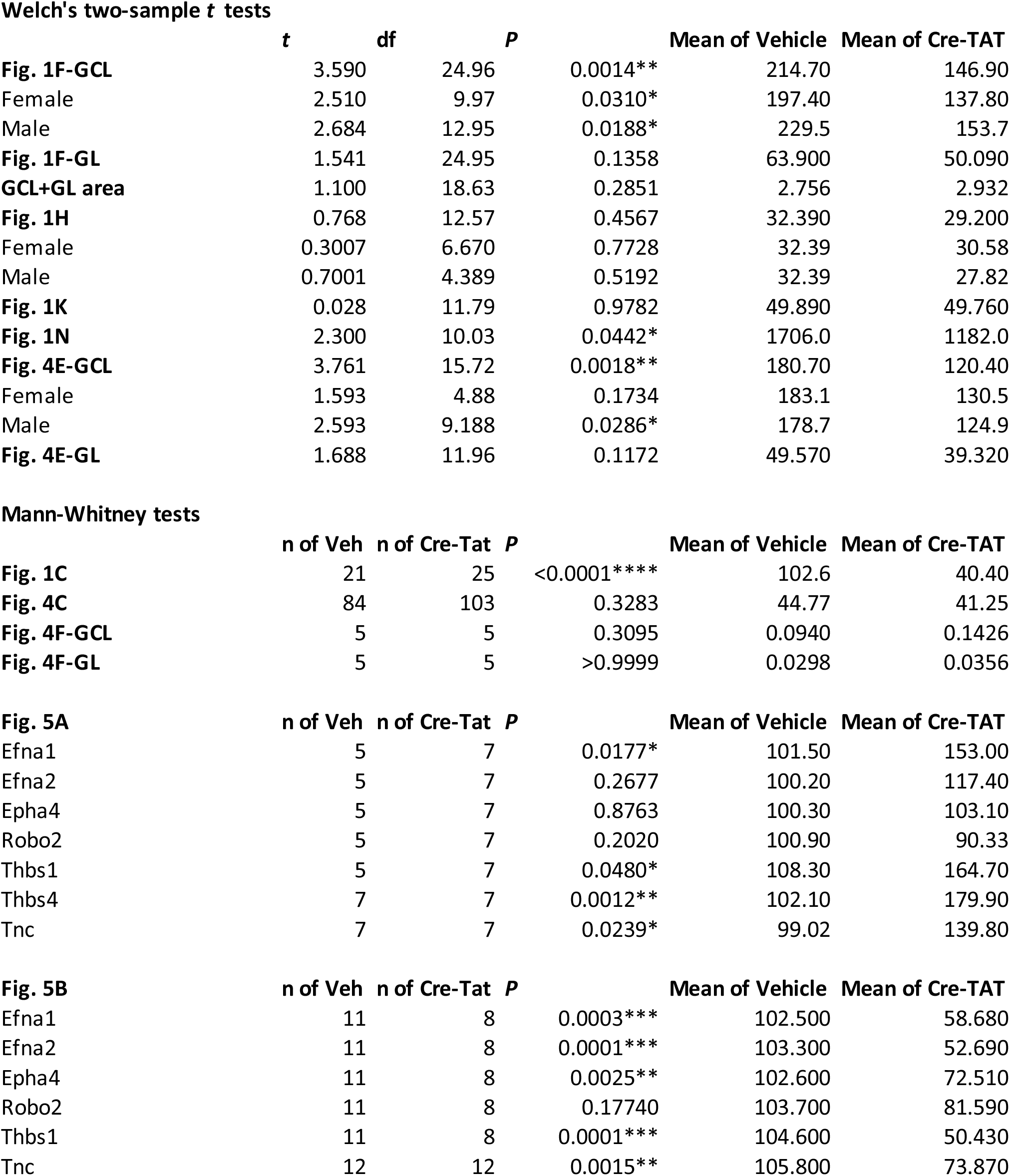
Statistical analysis.

### Non-cell-autonomous Otx2 in V-SVZ and RMS

Given that OTX2 is secreted from ChP into the CSF and accumulates in brain parenchyma (Spatazza et al., 2013), we hypothesized that OTX2 transfers from ChP to the V-SVZ niche. The *Otx2* locus is silent in V-SVZ and RMS, as shown by the absence of GFP expression in *Otx2^+/GFP^* mice (Fig. 2*A,B*) and by *Otx2 in situ* hybridization (Fig. 2*C*). However, OTX2 protein is immunoprecipitated from lysates of V-SVZ and RMS microdissections (Fig. 2*D*) but not from striatum lysates (microdissection beneath V-SVZ); lysates of superior colliculus, in which *Otx2* locus is active, were used as positive control (Fig. 2*E*). Immunostaining showed selective presence of OTX2 in a subset of cells within the V-SVZ and RMS (Fig. 3), suggesting it could be targeting either neural stem cells or progenitor cells directly and/or other cells within the niche. OTX2 is not internalized in neuroblasts (DCX+ cells) in V-SVZ but accumulates in a small number of GFAP-labeled cells and many ependymal cells (Fig. 3*A,B*). In and around the RMS, OTX2 is again absent in neuroblasts yet is in many GFAP labeled cells (Fig. 3*C,D*).

**Figure 3.**
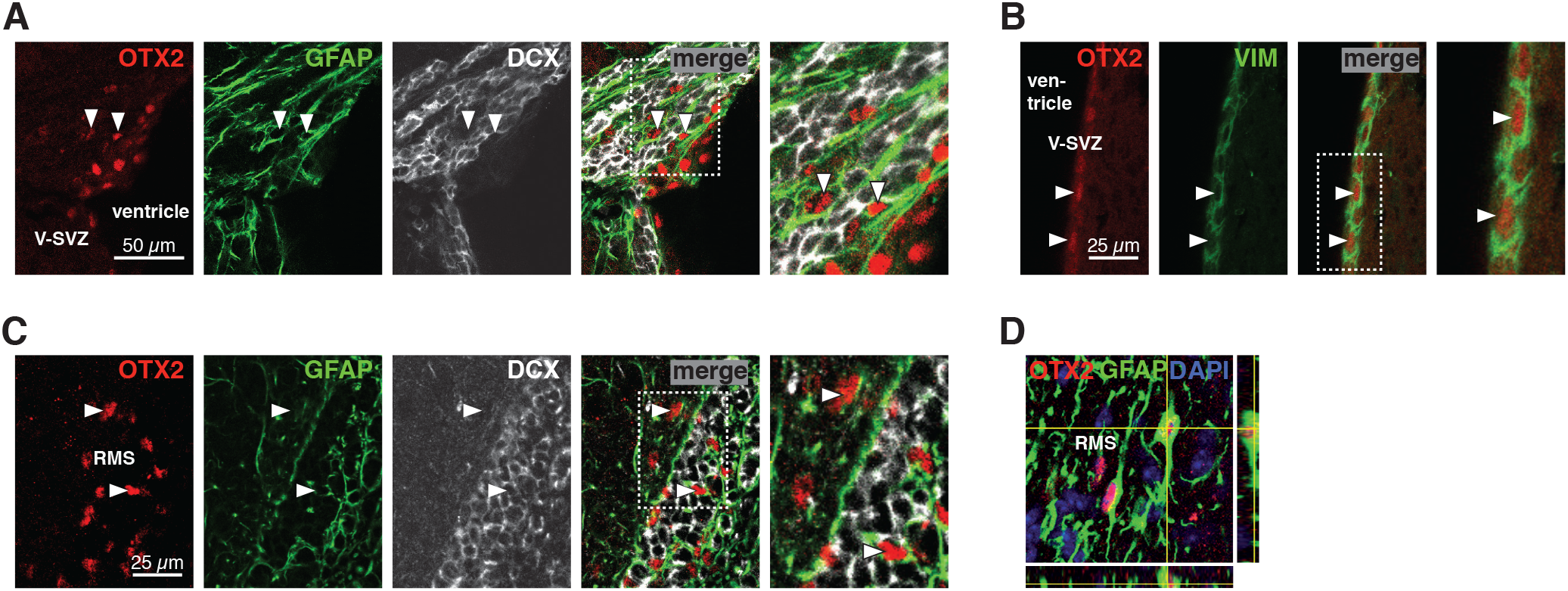
OTX2 transfers into supporting cells of neurogenic niches. ***A***, In dorsal V-SVZ, OTX2 staining is detected in some GFAP-labeled astrocytes but not in DCX-labeled neuroblasts. Arrowheads highlight astrocytes containing OTX2. ***B***, Along the lateral ventricular wall, OTX2 staining is detected in vimentin (VIM) labeled ependymal cells. ***C***, In the RMS, OTX2 staining is detected in GFAP-labeled astrocytes but not in DCX-labeled neuroblasts. Arrowheads highlight astrocytes containing OTX2. ***D***, Orthogonal projection shows OTX2 in the nucleus of astrocytes within the RMS. RMS, rostral migratory stream; V-SVZ, ventricular-subventricular zone.

OTX2 protein in regions of neuroblast proliferation and migration suggests that reduced newborn neurons in OB after *Otx2* ChP knockdown in Otx2-lox mice (Fig. 1*F*) could be due in part to OTX2 non-cell-autonomous activity. To test this hypothesis, we used the scFv-Otx2 mouse model which conditionally expresses a single chain antibody (scFv) that blocks transfer by sequestering OTX2 in the CSF (Bernard et al., 2016). Cre-Tat icv injection leads to local scFv-Otx2 expression by ChP cells (GFP reporter, Fig. 4*A*) and secretion into the CSF, which can lead to ~20% decrease in cortical OTX2 levels after 2 weeks (Bernard et al., 2016). Here, 35 days after Cre-Tat icv injection, this sequestering of Otx2 in CSF led to ~10% decrease in OTX2 levels within V-SVZ (Fig. 4*B,C*), although not statistically significant given the large distribution of OTX2 mean intensity per cell (Fig. 4*C*). Subjecting these mice to the BrdU paradigm for OB integration (Fig. 4*D*) revealed a decrease in newborn neurons in the GCL of Cre-Tat-injected animals and no significant change in the GL (Fig. 4*E*). TUNEL assay revealed no change in neuroblast survival upon arrival in the OB (Fig. 4*F*). Taken together, these results are similar to what was observed in Otx2-lox mice (Fig. 1*F,H*) and suggests that the transfer of OTX2 transfer from CSF to V-SVZ plays a significant role in regulating adult neurogenesis. The accumulation of OTX2 protein in supporting cells (Fig. 3) within the niche further suggests that OTX2 regulates V-SVZ neurogenesis though modification of the niche microenvironment.

**Figure 4.**
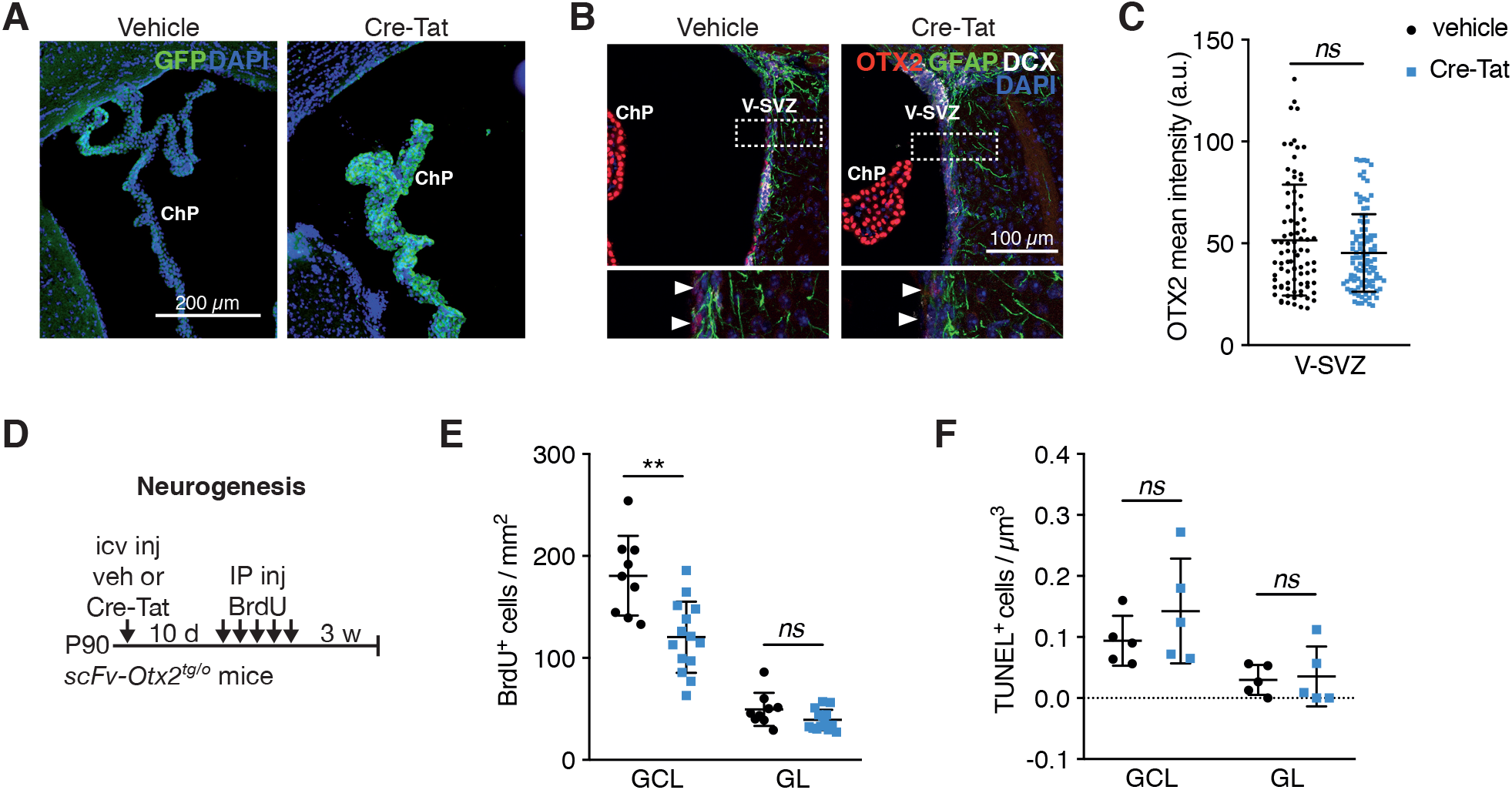
Non-cell autonomous OTX2 is sufficient to regulate adult neurogenesis. ***A***, Efficient recombination in choroid plexus of scFv-Otx2 mice after icv injection of Cre-Tat assessed by GFP reporter. ***B***, Change in OTX2 staining within V-SVZ 35 days after Cre-Tat icv injection. ***C***, Quantification of OTX2 mean intensity per cell in V-SVZ (*ns*=*P*>0.05, Mann-Whitney test). ***D***, Schematic of adult (3 months) neurogenesis study paradigm with scFv-Otx2 mice. ***E***, Quantification of BrdU-positive cells in olfactory bulb GCL and GL (*ns*=*P*>0.05, \*\**P*< 0.01, t-test, 3 experiments). ***F***, TUNEL analysis of GCL and GL to assess cell death in olfactory bulb of scFv-Otx2 mice (*ns*=*P*>0.05, Mann-Whitney test, 2 experiments). GCL, granule cell layer; GL, glomerular layer; icv, intracerebroventricular; V-SVZ, ventricular-subventricular zone. Error bars represent SD.

### OTX2 regulates astrocyte factors

As we observe OTX2 internalization all along the migrating route of neuroblasts (Fig. 3) and found decreased neuroblast migration (Fig. 1*N*) with no change in cell proliferation (Fig. 1*K*), we focused on a possible role for astrocytes found in both V-SVZ and RMS. Several receptors and factors are expressed in V-SVZ and RMS astrocytes that could potentially alter microenvironment composition and signaling. These include extracellular matrix (ECM) proteins such as thrombospondin (Thbs1 and Thbs4) and tenascin-C (Tnc) (Jankovski and Sotelo, 1996; Girard et al., 2014) and signaling molecules such as Robo and Eph receptors along with Ephrin ligands (Kaneko et al., 2010; Ming and Song, 2011; Falcão et al., 2012; Gengatharan et al., 2016; Todd et al., 2017), which may be under transcriptional control of OTX2 (Gherzi et al., 1997; Boncinelli and Morgan, 2001; Hoch et al., 2015; Peña et al., 2017). Expression analysis on V-SVZ microdissection lysates from the scFv-Otx2 non-cell autonomous OTX2 knock-down model showed that *EphrinA1, Thbs1, Thbs4* and *Tnc* were upregulated in V-SVZ (Fig. 5*A*). We also found that both *Thbs4* and *Tnc* were upregulated in V-SVZ of Cre-Tat injected Otx2-lox mice (304.6% ± 114.5 SD (*P*<0.0001) and 201.1% ± 94 SD (*P*=0.0037) of vehicle, respectively), reinforcing the similarity between both mice models.

**Figure 5.**
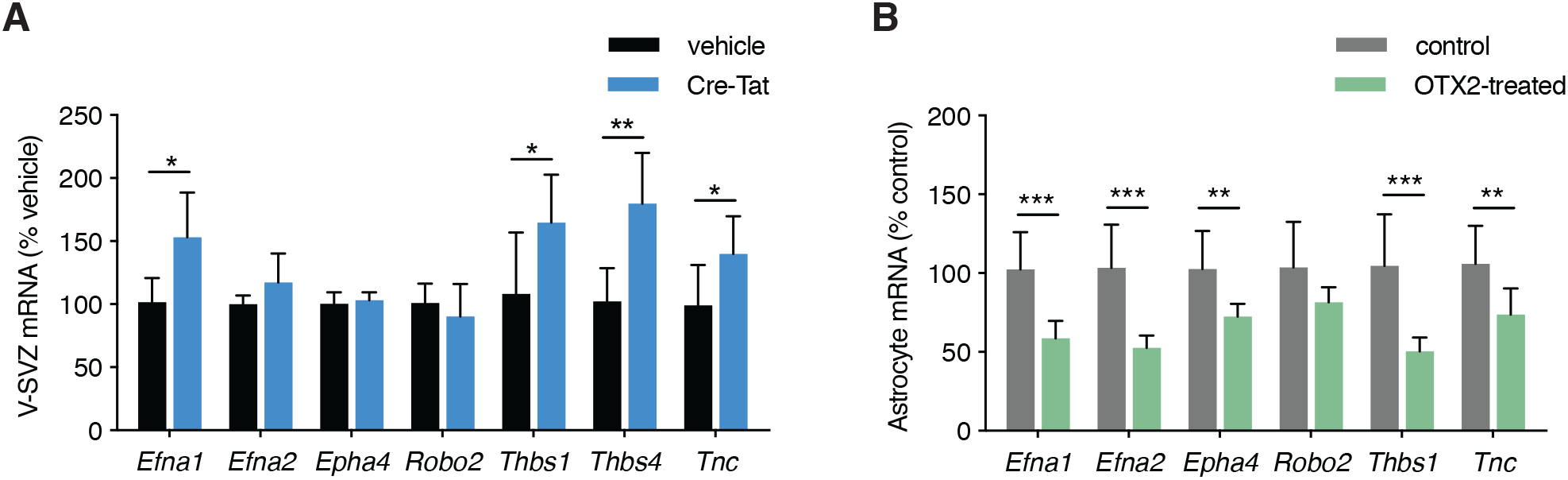
OTX2 regulates expression of ECM and signaling proteins. ***A***, Quantitative PCR analysis of V-SVZ gene expression in scFv-Otx2 after icv injection of Cre-Tat or vehicle (*ns*=*P*>0.05, \**P*<0.05, \*\**P*<0.01, Mann-Whitney test). ***B***, Quantitative PCR analysis of OTX2-treated astrocyte cultures (\*\**P*<0.01, \*\*\**P*< 0.001, Mann-Whitney test). V-SVZ, ventricular-subventricular zone. Error bars represent SD.

To confirm that supporting astrocytes are directly implicated in these changes in expression of ECM and signaling factors, we turned to an *in vitro* astrocyte model involving aged astrocytes from newborn mouse cortex to provide mature astrocytes that do not contain NSCs typical of V-SVZ cultures. Indeed, cultured late juvenile and adult cortex astrocytes do not give rise to neurons, while cultured V-SVZ astrocytes retain this potential (Laywell et al., 2000). However, astrocyte cultures derived from early juvenile mouse cortex show low *Thbs4* expression (Benner et al., 2013), and cultured astrocytes typically show enriched expression of *Thbs1* and repression of *Thbs4* compared to their *in vivo* counterparts (Cahoy et al., 2008; Eroglu, 2009). While astrocytes in our *in vitro* model do not perfectly represent isolated V-SVZ supporting cells, they are suitable for assessing non-cell autonomous OTX2 activity on mature and differentiated astrocytes given the absence of confounding NSCs. Consistent with changes observed *in vivo*, treatment with OTX2 protein resulted in the down-regulation of *EphrinA1, Thbs1* and *Tnc* (Fig. 5B), with the exception of *Thbs4*, which is not expressed in our cultured astrocytes and is in keeping with previous studies. In this *in vitro* paradigm, OTX2 was also found to regulate *EphrinA2* and *Eph4* expression which further highlights differences with adult V-SVZ astrocytes. Nonetheless, these results are in agreement with OTX2 transfer to niche astrocytes to regulate neurogenesis by controlling ECM composition and signaling.

## Discussion

OTX2 homeoprotein was previously found to act non-cell-autonomously in cortical interneurons (Sugiyama et al., 2008; Spatazza et al., 2013; Lee et al., 2017). The *Otx2* locus is not active in the V-SVZ and RMS, but the protein can be immunoprecipitated, indicating that OTX2 homeoprotein transfers not only into several distant cortical regions but also into regions proximal to the lateral ventricles. In the cortex, OTX2 is specifically internalized by parvalbumin interneurons, which have a unique and stable ECM called perineuronal nets (PNNs) that often stain for the lectin *Wisteria floribunda* (WFA) (Sugiyama et al., 2008). The specificity of internalization arises from a glycosaminoglycan-binding motif in OTX2 protein (Beurdeley et al., 2012; Lee et al., 2017), and although specific target glycosaminoglycans of OTX2 have yet to be identified, OTX2 preferentially binds di-sulfated chondroitin sulfate proteoglycans (Beurdeley et al., 2012; Miyata et al., 2012; Bernard and Prochiantz, 2016). However, staining for WFA is not detected in the V-SVZ (data not shown), which is not surprising given that PNNs are synonymous with consolidation of ECM and a break on plasticity - events that would counter the function of the neurogenic niche. The hypothesis of ECM-OTX2 interaction accounting for the specific internalization remains the most plausible, given that homeoprotein cell-surface receptors have not been found (Bernard and Prochiantz, 2016). Thus, the specificity of OTX2 for astrocytes within the V-SVZ and RMS suggests they are expressing glycosaminoglycans with high affinity for OTX2.

Through specific conditional recombination in the ChP of two mouse models, similar decreases in neurogenesis were observed. In Otx2-lox mice, recombination results in loss of local OTX2 expression in the ChP, while in scFv-Otx2 mice it results in sequestering OTX2 in the CSF due to single chain antibody secretion by the ChP (Bernard et al., 2016). The first model includes both potential cell-autonomous effects of loss of OTX2 in the ChP as well as potential non-cell-autonomous effects through reduced levels of secreted OTX2. The second model comprises only non-cell-autonomous effects as only secreted OTX2 is affected. This is confirmed by the absence of autocrine effects (i.e. CSF OTX2 signaling on ChP) assessed by ChP mRNA levels of OTX2 transcription targets upon scFv expression (Bernard et al., 2016). Since both models provide a similar reduction in the number of newborn neurons, it suggests that OTX2 non-cell-autonomous function is of primary importance for adult neurogenesis. Interestingly, we observe an effect despite a non-significant reduction in OTX2 accumulation in V-SVZ, which is likely due in part to the large distribution of mean OTX2 staining intensity. Small (~20%) decreases in OTX2 in adult mouse visual cortex induce cortical plasticity (Spatazza et al., 2013; Bernard et al., 2016), and OTX2 transcriptional targets can be activated at one concentration of OTX2 yet be suppressed at lower or higher concentrations (Apulei et al., 2018). Concentration-dependent targets of homeoprotein are shown to be dependent on chromatin state, whereby some targets are concentration insensitive while others are highly concentration-sensitive (Hannon et al., 2017). Indeed, we find significant changes in some but not all putative targets of OTX2 in the V-SVZ despite the potentially subtle changes in OTX2 accumulation.

Previous studies on the role of factors secreted by the ChP mainly show effects on the proliferation of NSCs and transit amplifying cells, as well as effects on maintaining NSC quiescence and the migration of neuroblasts (Bjornsson et al., 2015; Silva-Vargas et al., 2016). These factors were found to signal through quiescent and/or activated NSCs, such as for BMP5, IGF2 or NT-3 (Delgado et al., 2014; Ziegler et al., 2015; Silva-Vargas et al., 2016), or were found to signal directly through neuroblasts to regulate migration, such as for SLIT1/2 (Nguyen-Ba-Charvet et al., 2004). Interestingly, ROBO2 is expressed by supporting astrocytes within the V-SVZ and RMS and influences the migration of neuroblasts which express SLIT1 (Kaneko et al., 2010). This suggests that SLIT1/2 from ChP can also be signaling through supporting astrocytes. We find the ChP knockdown of OTX2 reduces GCL newborn neurons without significantly modifying of cell proliferation or cell death but by modifying neuroblast migration into the RMS. However, one might expect reduced neuroblast migration to lead to a proportional reduction in dying neuroblasts. Yet cell death is difficult to study in this population given the spatiotemporal heterogeneity of their death and the high variability in their detection. Thus, we cannot formally exclude that OTX2 is also affecting cell death. Nonetheless, OTX2 protein transfers in GFAP-positive cells in the V-SVZ, in the majority of GFAP-positive cells in the RMS, and in ependymal cells of the V-SVZ; this profile points to modification of the neurogenic niche instead of direct modification of progenitor activity.

Given the precedent of ECM and supporting astrocytes for regulating neuroblast migration (Gengatharan et al., 2016; Kaneko et al., 2017), we examined OTX2-dependent expression of local ECM and signaling molecules expressed within V-SVZ and RMS. The RMS is composed of a compacted neuroblast network forming chains that migrate along blood vessels and are surrounded by astrocytic processes (Lois et al., 1996; Whitman et al., 2009). While it is clear that RMS-associated astrocytes interact with neuroblasts (Kaneko et al., 2010), the role of astrocytes for neuroblast migration is not well known. Studies show that neurogenesis is impacted by the composition of ECM in stem cells niches (Dityatev et al., 2010), which include astrocyte-derived TNC and THBS4 in V-SVZ and RMS (Jankovski and Sotelo, 1996; Benner et al., 2013; Girard et al., 2014), and OTX2 has been shown to regulate *Tnc* and *Thbs4* transcription (Gherzi et al., 1997; Peña et al., 2017). While *Tnc* knockout mice show no signs of altered adult neurogenesis (Kazanis et al., 2007), *Thbs4* knockout adult mice show reduced newborn neurons in the OB yet no defect of embryonic neurogenesis (Girard et al., 2014). Cell-autonomous OTX2 expression was found to parallel *Thbs4* expression in the retina (Housset et al., 2013), suggesting that OTX2 activity is context dependent as previously suggested (Buecker et al., 2014). However, given the global dysregulation observed, we cannot conclude on the role of the different ECM molecules and interacting proteins. Nevertheless, changes in V-SVZ expression of *Tnc* and *Thbs4* upon *Otx2* knockdown are in-keeping with a transcriptional role for non-cell autonomous OTX2 in astrocytes.

As OTX2 was found in RMS astrocytes and that signalization with neuroblasts is still an open question (Chaker et al., 2016), we tested if genes coding for migration and signaling factors might be modified in our models. *Robo2* is strongly expressed in RMS astrocytes and is required for proper RMS organization and guidance of neuroblasts (Kaneko et al., 2010), and it has been hypothesized that astrocyte-derived laminin helps form RMS scaffold for neuroblast migration (Belvindrah et al., 2007). Also, *EphA4* and *EphrinA2* are expressed in RMS and V-SVZ supporting astrocytes (along with other combinations of Ephs and Ephrins) and EPHA4 has been shown to be critical for maintaining a compact and organized RMS (Todd et al., 2017). While OTX2 has been shown to bind *Robo2* promoter (Hoch et al., 2015) and to regulate *Ephrin-A2* (Rhinn et al., 1999), we did not detect any changes in V-SVZ expression after OTX2 knock-down. However, OTX2 affected expression more widely in cultured mature astrocytes, including expression of *EphA4* and *EphrinA2*, suggesting it has the potential to influence RMS organization and/or that its activity is differentially regulated among astrocyte subtypes. While these cultured astrocytes are not derived from V-SVZ, they reflect better the activity of OTX2 in supporting astrocytes given that V-SVZ cultures are dominated by GFAP-expressing NSCs. This *in vitro* model confirms that transfer of OTX2 into V-SVZ can regulate the expression of genes found in supporting astrocytes.

Our results add to the growing research on homeoprotein transduction (Di Nardo et al., 2018) and add to the complexity of ChP function, which is essential for brain homeostasis and neuroimmune regulation (Spector et al., 2015; Ghersi-Egea et al., 2018). OTX2 signaling through the CSF to cortical interneurons has been shown to regulate brain plasticity throughout the cortex and impact higher-cognitive functions. We must now consider that ChP-derived OTX2 accumulates in diverse regions of the brain and that altering its concentration within CSF can influence a wide range of brain functions. Within the V-SVZ, OTX2 regulates ECM molecules and factors that likely impact neuroblast migration and ultimately newborn neuron numbers. These findings reinforce both ChP and V-SVZ non-neurogenic astrocytes as key niche compartments affecting adult neurogenesis.

Author Contributions
APl, VOM, APr and AAD designed the experiments; APl, VOM and CD performed the experiments; APl, VOM and AAD wrote the manuscript.

